# DeepPerVar: a multimodal deep learning framework for functional interpretation of genetic variants in personal genome

**DOI:** 10.1101/2022.04.10.487809

**Authors:** Ye Wang, Li Chen

**Affiliations:** Biotherapeutic and Medicinal Sciences, Biogen, 225 Binney Street, Cambridge, MA, 02142, USA; Department of Biostatistics and Health Data Science, Indiana University School of Medicine, Indianapolis, IN, 46033; Center for Computational Biology and Bioinformatics, Indiana University School of Medicine, Indianapolis, IN, 46033

## Abstract

**Motivation:** Understanding the functional consequence of genetic variants, especially the noncoding ones, is important but particularly challenging. Genome-wide association studies or quantitative trait locus analyses may be subject to limited statistical power and linkage disequilibrium, and thus are less optimal to pinpoint the causal variants. Moreover, most existing machine learning approaches, which exploit the functional annotations to interpret and prioritize putative causal variants, cannot accommodate the heterogeneity of personal genetic variations and traits in a population study, targeting a specific disease.

**Results:** By leveraging paired whole genome sequencing data and epigenetic functional assays in a population study, we propose a multi-modal deep learning framework to predict genome-wide quantitative epigenetic signals by considering both personal genetic variations and traits. The proposed approach can further evaluate the functional consequence of noncoding variants on an individual level by quantifying the allelic difference of predicted epigenetic signals. By applying the approach to the ROSMAP cohort studying Alzheimer’s disease (AD), we demonstrate that the proposed approach can accurately predict quantitative genome-wide epigenetic signals and in key genomic regions of AD causal genes, learn canonical motifs reported to regulate gene expression of AD causal genes, improve the partitioning heritability analysis, and prioritize putative causal variants in a GWAS risk locus. Finally, we release the proposed deep learning model as a stand-alone Python toolkit and a web server.

**Availability:** https://github.com/lichen-lab/DeepPerVar

## 1 Introduction

Studying the functional consequences of genetic variants is an important research area of genetics. Interpreting the functional consequences of protein coding variants has been successful, which has improved the understanding of the disease biology and drug discovery [1]. However, deciphering functional roles of noncoding variants in regulatory DNA is still challenging. Genome-wide association studies (GWAS) have been widely used to identify disease-associated risk variants, most of which fall within the noncoding regions [2]. However, GWAS have so far remained largely underpowered in identifying associations of the rare variants and have lacked the resolution to pinpoint causal variants due to linkage disequilibrium [3, 4, 5]. The advent and popularity of whole genome sequencing data (WGS) makes the identification of rare variants possible. Understanding rare and individual-specific variants will not only explain the missing heritability from GWAS but also improve the precision medicine. However, it is still a challenging to identify disease-associated rare variants due to the limited sample size of expensive sequencing data. Statistical methods such SKAT [6] has been developed to ameliorate this issue by performing the SNP-set (e.g., SNPs in a gene or a region) wise test instead of SNP-wise test, which significantly improve the statistical power. However, the association test is on SNP-set, so it still is difficult to pinpoint which SNP is causal in the SNP set. In addition, both identified common and rare variants by either GWAS or WGS analysis utilize genotype data only. However, there is a lack of functional insights on how these variants affect molecular mechanisms on the tissue and cell type level in order to have an impact on the disease.

To provide the biological insights to interpret the noncoding variants, tissue and cell type-specific functional annotations, which are mainly derived from public consortiums such as ENCODE [7] and Roadmap Epigenomics Project [8], have been integrated with GWAS/WGS summary statistics to enable the refinement of disease risk associated region and prioritization of causal variants. The first attempt towards this direction is to perform the positional overlap between GWAS SNPs and regulatory regions [9, 10, 8, 11] by assuming that the SNP is more likely to be casual if it is more enriched with functional annotations. Moreover, multiple machine learning models have recently been developed to leverage labelled known causal variants in Human Gene Mutation Database (HGMD) [12] and ClinVar [13], and multi-omics functional annotations. These methods achieve a genome-wide prediction by assigning a functional score to each variant [14, 15, 16, 17, 18, 19, 20, 21, 22]. However, despite the success of these annotation-based approaches, the allelic effect of a variant cannot be assessed.

With the advance of deep learning research in the genomic area, deep convolutional neural network (CNN), have been developed to predict the regulatory activity in epigenetic regions using DNA genomic sequence. By comparing the prediction using reference allele and alternative allele in the DNA sequence, these methods can *in silico* evaluate allelic effect of a variant by taking the difference of the predicted regulatory activity. State-of-the-art method, such as DeepSEA [23] and its extension ExPecto [24], is a multi-task CNN to provide a binary quantification (e.g. ChIP-seq peak region or not) of genome-wide histone modification, transcription factor binding and chromatin accessibility across multiple cell types and tissues simultaneously. However, only a large effect size of epigenetic signal can result in a peak region, but it is still crucial to evaluate the functional consequence of a variant located in a genomic region of moderate or small effect size of epigenetic signal, which is not large enough to generate a peak. Moreover, real applications such as epigenetic imputation using DNA sequence requires the quantitative prediction instead of qualitative prediction. These considerations make the quantitative prediction more favorable. Moreover, these cutting-edge methods use reference genome sequences as model input, making them are less optimal to model the contribution of genetic variations driving the epigenetic signal. However, the heterogeneity of genetic variations, especially in a population study, should be considered in the model training.

Large public consortiums provide paired genotype data (e.g., SNP array, WGS) and molecular phenotype data (e.g., RNA-seq, ChIP-seq, ATAC-seq, DNA methylation data) in a population study to study different disease etiology. For example, TCGA [25] studies different cancer types; PsychENCODE [26] focus on autism spectrum disorder, bipolar disorder, and schizophrenia; and ROSMAP [27] targets Alzheimer’s disease. These resources provide an unprecedented opportunity to evaluate and interpret the functional consequence of genetic variants by integrating molecular phenotypes. A common practice for integrating genotype data and molecular phenotype data is Quantitative trait locus (QTL) analysis. However, QTL analysis faces the same challenge as GWAS such as linkage disequilibrium and statistical power. A recent developed deep learning method, named DeepFIGV, integrates paired WGS and ChIP-seq data to predict quantitative epigenetic variation in chromatin accessibility and histone modifications in lymphoblastoid cell lines across 75 individuals [28]. Though DeepFIGV adopts personal genetic variations in the model training, it did not consider personal traits, such as age and gender, as well as the interaction between genetics and traits, which are important factors associated with epigenetic change.

Here, we introduce DeepPerVar, which is a deep learning framework to perform the functional interpretation of genetic variants in personal genome by leveraging paired whole genome sequencing data and epigenetic functional assays in a population study. DeepPerVar is essentially a multi-modal deep convolutional neural network, which considers both personal genome and personal traits, as well as their interactions in the model training, to quantitatively predict epigenetic signals and evaluate the functional consequence of genetic variants on an individual level. To demonstrate its usefulness, we apply DeepPerVar to ROSMAP, which is an AD study cohort with paired WGS data and H3K9ac ChIP-seq data as well as DNA methylation data. We demonstrate that DeepPerVar achieves an accurate prediction for genome-wide epigenetic signals and in key genomic regions associated with AD, discover canonical motifs and motifs associated with AD and prioritize putative causal variants associated with AD in a GWAS locus. Especially, DeepPerVar outperforms state-of-the-art methods DeepFIGV and ExPecto in partitioning heritability analysis. Finally, DeepPerVar is released as both a stand-alone command line tool and a web server, which can provide the functional score given the genomic loci of genetic variants. We believe DeepPerVar will be a useful tool for interpreting rare and individual-level genetic variants in a population study.

## 2 Methods

### 2.1 Paired whole genome sequencing and epigenetic datasets from ROSMAP

The whole genome sequencing data (WGS) and epigenetic data, which include H3K9ac ChIP-seq data and 450K DNA methylation array derived from dorsolateral prefrontal cortex, as well as phenotypic data of individuals are collected from the “ROSMAP” study [27]. In the ROSMAP study, we select the “Definite” AD patients using CERAD score with value 1. As a result, we obtain 202 AD patients profiled with both WGS data and 450K DNA methylation arrays, and 196 AD patients profiled with both WGS data and H3K9ac ChIP-seq data.

The WGS data in the format of genomic VCF files are generated by GATK [45], which adopts a rigorous quality assessment for variant filtration to produce detected germline WGS single nucleotide variants (SNVs) mapped to GRCh37/hg19 reference genome. For H3K9ac ChIP-seq data, we identify ChIP-seq peaks from bam files using MACS2 [46] with additional two quality control criteria to remove low-quality H3K9ac peaks: i) ChIP counts are smaller than matched input control counts; ii) p-value is less than 0.05 derived from the Poisson test with normalized ChIP counts as observed value and smoothing input control counts as Poisson parameter. We merge overlapped peaks across all individuals and calculate the normalized read counts for each merged peak by adjusting sequence depth and matched input control. Eventually, we have 141,807 merged peaks for downstream analysis.

The 450K DNA methylation data have been imputed and undergo a series of quality control analysis and have been adjusted for age, sex, and experimental batch, which ends up with methylation ratio at 418,972 CpGs [27]. We then use R/Bioconductor package “IlluminaHumanMethylation450kanno.ilmn12.hg19” to obtain the genomic coordinates for all CpGs and annotate CpGs in four methylation regions, which results in 135,303 CpGs in island, 98,617 CpGs in shore, 39,144 CpGs in shelf, and 145,938 CpGs in open sea.

### 2.2 Epigenetic datasets from Roadmap Epigenomics Project

We collect epigenetic datasets from multiple tissues and cell lines in human from the Roadmap Epigenomics Project [8]. For ChIP-seq data, we collect a total of 439 datasets, which are across 83 cell lines/tissues, 18 tissue classes and 7 core histone marks including H3K4me1, H3K9ac, H3K4me3, H3K9me3, H3K27ac, H3K27me3 and H3K36me3. For DNA methylation data, we obtain 104 DNA methylation datasets generated from three different sequencing technologies, which include Reduced representation bisulfite sequencing (RRBS), Whole-Genome Bisulfite Sequencing (WGBS), and mCRF across multiple cell lines. Specifically, RRBS data consists of 51 datasets across 51 cell lines/tissues and 13 tissue classes. WGBS data consists of 37 datasets across 37 cell lines/tissues and 13 tissue classes. mCRF data consists of 16 datasets across 16 cell lines/tissues and 5 tissue classes. All data have been preprocessed following the Roadmap protocol, which provides the methylation ratio for each CpG and normalized ChIP read counts for each peak.

### 2.3 Constructing personal genome sequence and personal trait

We define the epigenetic region as flanking region of ChIP-seq peak or CpG site. Without loss of generality, we extend 500bp upstream and downstream of each CpG site or 1000bp upstream and downstream of center of each ChIP-seq peak to allow the same dimensionality of model input for the neural network. Next, we adopt bcftools tools [47] to obtain the reference genome sequence from the epigenetic region, and edit the sequence by incorporating biallelic SNPs from the WGS VCF files to construct personal genome sequence. Specifically, homozygous alternate site can be directly replaced by the alternative allele. Heterozygous site can be represented by the IUPAC nucleotide codes because one reference allele can mutate into different types of alternative alleles. For example, heterozygous site “A/G” can be represented by IUPAC nucleotide code “R” in the DNA sequence. Afterwards, the personal genome sequence is further converted into one-hot encoded matrix, where homozygous reference/alternate site is represented as “A”: [1,0,0,0], “C”: [0,1,0,0], “G”: [0,0,1,0], “T”: [0,0,0,1], and heterozygous site such as “R” is represented as “A/G”: [0.5,0,0.5,0].

Moreover, we collect key phenotypic/clinic features, which include age of death, gender, years of education, APOE genotype, CREAD score, Braak staging and clinical cognitive diagnosis. Details of the distribution of the phenotypic data can be found in Supplementary Table S1. We further apply minimax-normalization to normalize these features.

### 2.4 Deep convolutional neural network architecture of DeepPerVar

DeepPerVar is essentially a multi-modal deep convolutional neural network, which consists of two modalities. The first modality/subnetwork is a hybrid of CNN-LSTM network, which aims to extract high-level feature representations of personal genome sequence in the epigenetic regions. The second modality/subnetwork adopts a feed-forward neural network to generate feature representations for personal traits. Finally, a tensor fusion layer models the unimodal representation and bimodal interactions of feature maps from both subnetworks and output the final feature map for prediction.

We denote the sequence subnetwork *𝒰*_*g*_, which learns a rich representation of the personal genomic sequence (Figure 2). Let **x** = {*x*_1_, *x*_2_, *x*_3_, ⋯, *x*_*m*_} as be the one-hot encoding representation of personal genomic sequence, where *m* is the length of the sequence. Then, a convolution layer acts as a motif scanner on **x** to produce a feature map *X*_*t*_ for each convolution kernel *t*. A Bidirectional LSTM layer with a forget gate is employed to learn the oriented and spatial dependency among motif representations, which is formularized as,

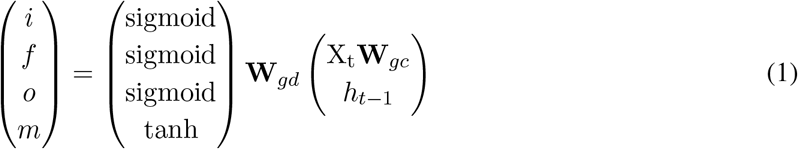

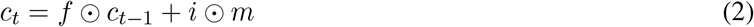

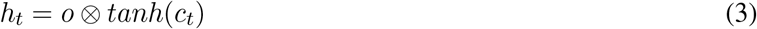

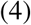

where *i, f, o, m, h*_*t*_, *c*_*t*_ are the input gate, forget gate, output gate, candidate cell state, hidden state and cell state, respectively; **W**_*gd*_ and **W**_*gc*_ are the weights of the Bidirectional LSTM layer. Two feature maps 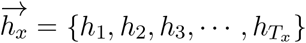 and 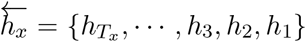 are generated after the Bidirectional LSTM layer is operated in the and forward and backward direction. Finally, a feature map 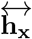, which is the output from the Bidirectional LSTM layer, is defined by concatenating both feature maps in the way as 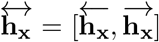. 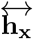 further connects a feed-forward neural network *f*_*g*_ to generates the feature map **z**^*x*^ from *𝒰*_*g*_, denoted as,

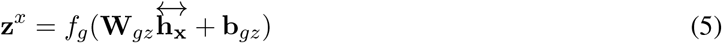

where **W**_*z*_ and **b**_*z*_ are the weight and bias for the feed-forward neural network. In short, **z**^*x*^ can be written as,

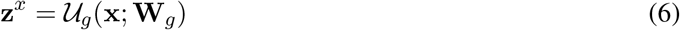

where **W**_*g*_ is the set of all weights in the *𝒰*_*g*_ network.

We denote the trait subnetwork as *𝒰*_*c*_, which is based on feed-forward neural network, to learn feature representations for personal traits. Let the personal traits as **v** = {*v*_1_, *v*_2_, *v*_3_, ⋯, *v*_*p*_*}. 𝒰*_*c*_ will output the feature map **z**^*v*^,

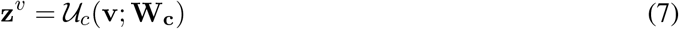

To integrate feature map **z**^*x*^ from trait subnetwork *𝒰*_*c*_, and **z**^*v*^ from sequence subnetwork *𝒰*_*g*_, we develop a fusion layer that disentangles unimodal and bimodal dynamics by modeling each of them explicitly, which is defined by the Cartesian product,

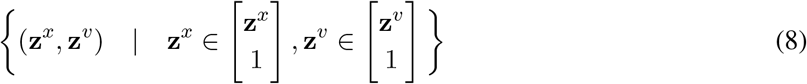

The extra constant scalar with value 1 is used to generate the unimodal and bimodal dynamics. This definition is equivalent to a differentiable outer product between **z**^*x*^ and **z**^*v*^,

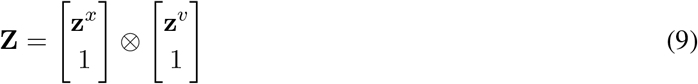

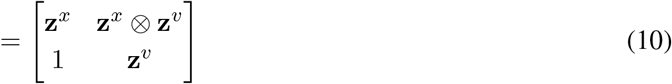

where ⊗ indicates the outer product between vectors. **Z** is a 2D matrix contains both unimodal and bimodal representations. Specifically, **z**^*x*^ and **z**^*v*^ are unimodal representations from sequence and trait subnetwork respectively. **z**^*x*^ ⊗ **z**^*v*^ captures bimodal interactions between **z**^*x*^ and **z**^*v*^ in tensor fusion layer. After the multimodal fusion, the **Z** will be flatten into **z** and fed to another feed-forward neural network *f*_*p*_ for prediction,

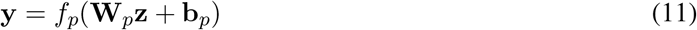

### 2.5 Model training, testing and software implementation

DeepPerVar is trained using the Adam algorithm [49] with a minibatch size of 256 to minimize the mean square error in the training set. Validation loss is evaluated at the end of each training epoch to monitor convergence. The weights of convolutional and dense layers are initialized by randomly Xavier uniform distribution. The orthogonal initialization is used to initialize the weights in Bidirectional LSTM layers. To improve generalization and to prevent overfitting, we use both *L*_2_ regularization for the weights and dropout [50]. Specifically, the dropout rates are set as 50% for both convolutional layer and dense layer.

To perform a conservative evaluation of the prediction performance, we conduct a “cross-sample cross-chromosome” approach to design training, validation, and testing sets. As a result, the epigenetic regions in training, validation, and testing sets are from different chromosomes in different individuals. Specifically, epigenetic regions in chromosome 1-8 from 60% individuals to the training set, epigenetic regions on chromosome 16-22 from 20% individuals to the validation set for hyper-parameter tuning and model selection, and epigenetic regions on chromosome 9-15 from the remaining 20% individuals are held-out for independent testing.

DeepPerVar stand-alone python toolkit is implemented by Pytorch [48] and the web server is developed using Google Cloud Platform. Given the genomic coordinate on hg19/GRCh37 of a variant, the predicted epigenetic signals such as H3K9ac signals and DNA methylation ratios, as well as the functional score, which is the allelic difference of the predicted epigenetic signals, will be reported and downloaded.

### 2.6 Variant effect prediction

Given a sequence variant, DeepPerVar (*ℱ*) will predict the quantitative epigenetic signals for both alternative and reference alleles. The functional score *δ* is defined as the absolute value of the difference,

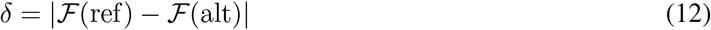

### 2.7 Comparison to canonical motifs

We collect canonical motifs of Homo Sapiens transcription factors in the format of position weight matrix (PWM) from CIS-BP Database [51] and motifs of transcription factors reported to be associated with Alzheimer’s disease [32]. To obtain the *de novo* motifs, we utilize all convolutional filters in the first layer of the DeepPerVar. To compare *de novo* motifs to the canonical motifs, we adopt Tomtom [52] and visualize the motifs using ggseqlogo [53].

## 3 Results

### 3.1 Both personal genetic variations and traits are associated with epigenetic signals

Most existing deep learning methods use DNA sequence from reference genome to predict epigenetic signals, and therefore do not consider the personal genetic variations within the sequences, which are widely present in a population study [23, 24]. In addition, personal traits such as gender and age are important factors associated with the change of epigenetic signals [30]. Essentially, it is the interaction between personal traits and genetic landscape that shape the epigenetic signals. To demonstrate this, we obtain the WGS genotype data and H3K9ac ChIP-seq data for AD patients from ROSMAP. First, we exam three male patients with one biallelic variant in two intronic regions of UGDH, STPG1 and one intergenic region (Figure 1A). For each region, we find that there is an additive effect, where the H3K9ac signals decrease as the genotype of one variant changes from homozygous reference alleles to heterozygous alleles to homozygous alternative alleles. Similarly, in another two intronic regions of DD39B, RAF1 and one intergenic region, we find H3K9ac signals increase as the genotype of one variant changes from homozygous reference alleles to homozygous alternative alleles (Figure 1B). This observation demonstrates that genetic variations are correlated with epigenetic signals. Second, we evaluate the H3K9ac signals for two men and two women in two intronic regions of ZNF622, WASH7P and one intergenic region (Figure 1C). Each region has one biallelic variant with the same genotype across four patients, which contains either homozygous reference/alternative alleles or heterozygous alleles. We find that H3K9ac signals of males are consistently higher than females in both intronic and intergenic regions for either homozygous or heterozygous alleles, which indicate that epigenetic signals are associated with gender. Overall, both findings confirm that both personal genetic variations and traits are associated with epigenetic signals and should be jointly considered in predictive modeling of epigenetic signals.

**Figure 1:**
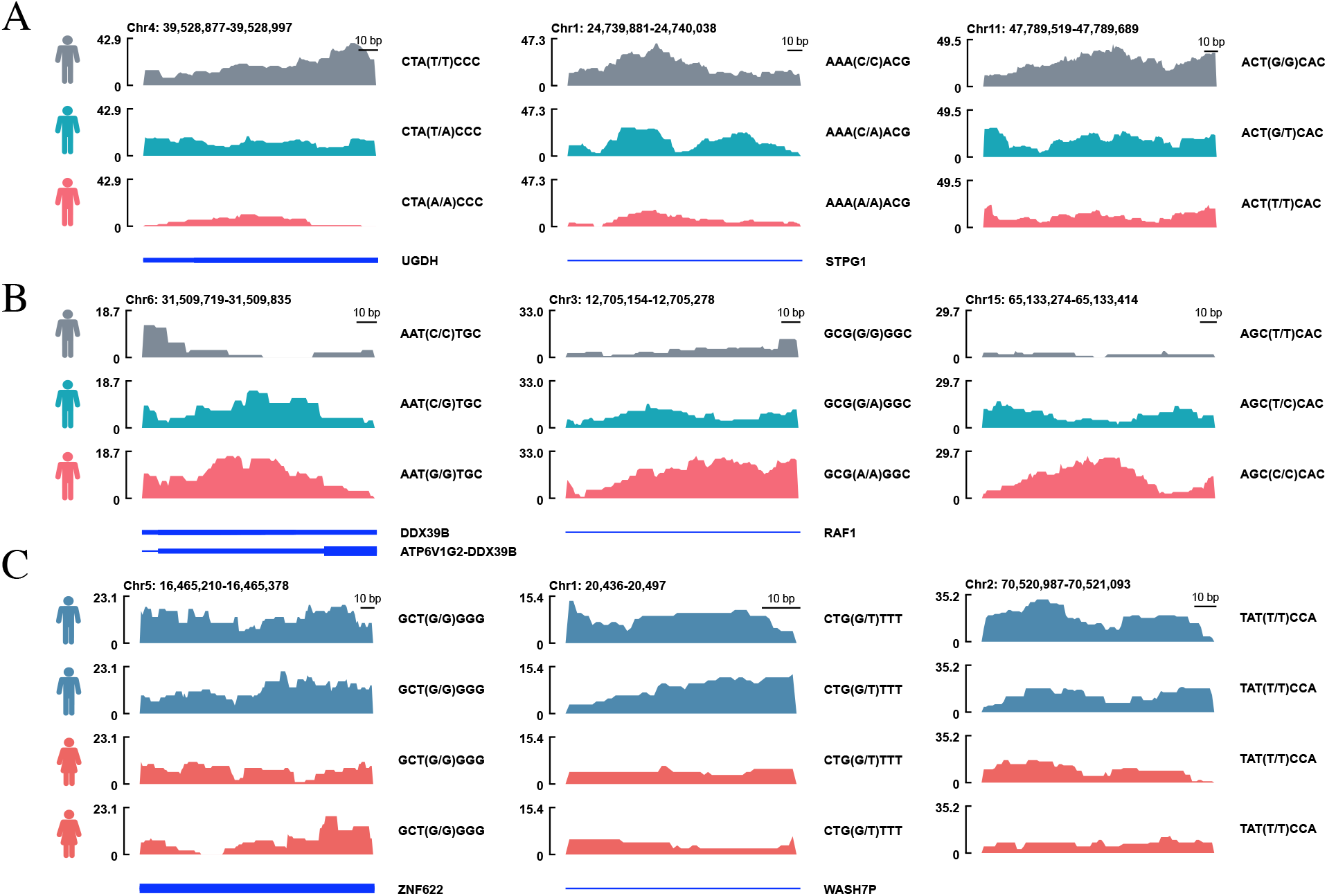
Both personal genetic variations and traits are associated with epigenetic signals. (A) For three male patients, the H3K9ac signals decrease as the genotype of one biallelic variant changes from homozygous reference alleles to heterozygous alleles to homozygous alternative alleles in two intronic regions of UGDH, STPG1 and one intergenic region. (B) For three male patients, the H3K9ac signals decrease as the genotype of one biallelic variant changes from homozygous reference alleles to heterozygous alleles to homozygous alternative alleles in two intronic regions of DD39B, RAF1 and one intergenic region. (C) For two males (marked blue) and two females (marked red) in two intronic and one intergenic region. Males have higher H3K9ac signals than females given the same genotype containing either homozygous or heterozygous alleles.

### 3.2 Deep convolutional neural network architecture of DeepPerVar

DeepPerVar is essentially a multi-modal deep convolutional neural network, which consists of two modalities. The first modality/subnetwork is a hybrid of CNN-LSTM network, which aims to extract high-level feature representations of personal genome sequence in the epigenetic regions. The epigenetic regions can be the flanking regions of methylation sites or histone modification sites. Personal genome sequence is created by obtaining reference genome sequence from the epigenetic regions first and then being edited to include personal genetic variants. Afterwards, one-hot encoding personal genome sequence will serve as the modality input. The connected convolutional layer will perform high-order feature extraction, followed by one Bidirectional LSTM layer to learn the long distance dependency of extracted features. The feature map output from the Bidirectional LSTM layers are further flattened and connect to the dense layers, which will generate final feature representations for personal genome sequence. The second modality/subnetwork adopts feedforward neural network to generate feature representations for personal traits. To integrate feature representations from both personal genome sequence and personal traits, DeepPerVar also adopts a tensor fusion layer to model the unimodal feature representations and bimodal interactions using a Cartesian product, which will generate an integrative feature map to dense layers for predicting quantitative epigenetic signals.

### 3.3 Predicting quantitative epigenetic signals by integrating both personal genomic sequence and trait

To demonstrate DeepPerVar can predict epigenetic signals accurately, we collect 196 paired whole genome sequencing (WGS) and DNA methylation array, and 202 WGS and H3K9ac ChIP-seq data from individuals diagnosed with “definite” AD in Religious Orders Study/Memory and Aging Project (ROSMAP) [27]. We perform a series of data processing steps to obtain epigenetic regions, which are flanking regions of ChIP-seq peaks and CpG sites. The CpG sites can be further classified into four regions, which include CpG island, shelf, shore and open sea regions (See Method section). By extending the center of ChIP-seq peak upstream and downstream 1000bp, and CpG site upstream and downstream 500bp, we obtain a fixed length of 2000bp reference genome sequence for ChIP-seq peak and 1000bp reference genome sequence for CpG site respectively. To create personal genome sequence, we further edit the reference genome sequence by including biallelic variants in the epigenetic region from a personal genome. Since each sequence comes from one individual, we further include personal traits such as gender, education level, and APOE genotype as the model input. The normalized read count in ChIP-seq peak and methylation ratio of CpG site are treated as the quantitative epigenetic signals. Finally, we train two versions of DeepPerVar, which are named “DeepPerVar-H3K9ac” and “DeepPerVar-methy” to predict normalized read count of H3K9ac and methylation ratio of CpG site respectively.

To perform a conservative evaluation of the prediction performance, we conduct a “cross-individual cross-loci” approach to design training, validation, and testing sets. Specifically, epigenetic regions in chromosome 1-8 from 60% individuals to the training set, epigenetic regions on chromosome 16-22 from 20% individuals to the validation set for hyper-parameter tuning and model selection, and epigenetic regions on chromosome 9-15 from the remaining 20% individuals are held-out for independent testing. As a result, the personal sequences in training, validation, and testing sets are from different individuals and different chromosomes. Since the outcome is quantitative signal, we adopt mean square error as the loss function for training the model. To measure the prediction performance, we calculate Pearson correlation (*R*) and Spearman’s rank correlation coefficient *ρ* between predicted and observed epigenetic signals.

Consequently, we find that *R* ranges from 0.677 to 0.713 (sd=0.013) and *ρ* ranges from 0.475 to 0.520 (sd=0.016) to predict H3K9ac signals (Figure 3A). Moreover, we find that the prediction accuracy for DNA methylation ratios varies across different methylation regions: *R* ranges from 0.691 to 0.757 (sd=0.024) and *ρ* ranges from 0.593 to 0.617 (sd=0.014) for CpG island; *R* ranges from 0.663 to 0.718 (sd=0.018) and *ρ* ranges from 0.687 to 0.724 (sd=0.015) for shore; *R* ranges from 0.275 to 0.335 (sd=0.022) and *ρ* ranges from 0.183 to 0.254 (sd=0.029) for shelf; *R* ranges from 0.397 to 0.500 (sd=0.043) and *ρ* ranges from 0.283 to 0.370 (sd=0.032) for open sea. Clearly, the prediction for CpG island and shore are more accurate than shelf and open sea, which may be explained by a higher density of CpGs in island and shore compared to shelf and open sea. The above results show that DeepPerVar can accurately predict quantitative epigenetic signals for both histone modification and DNA methylation.

**Figure 2:**
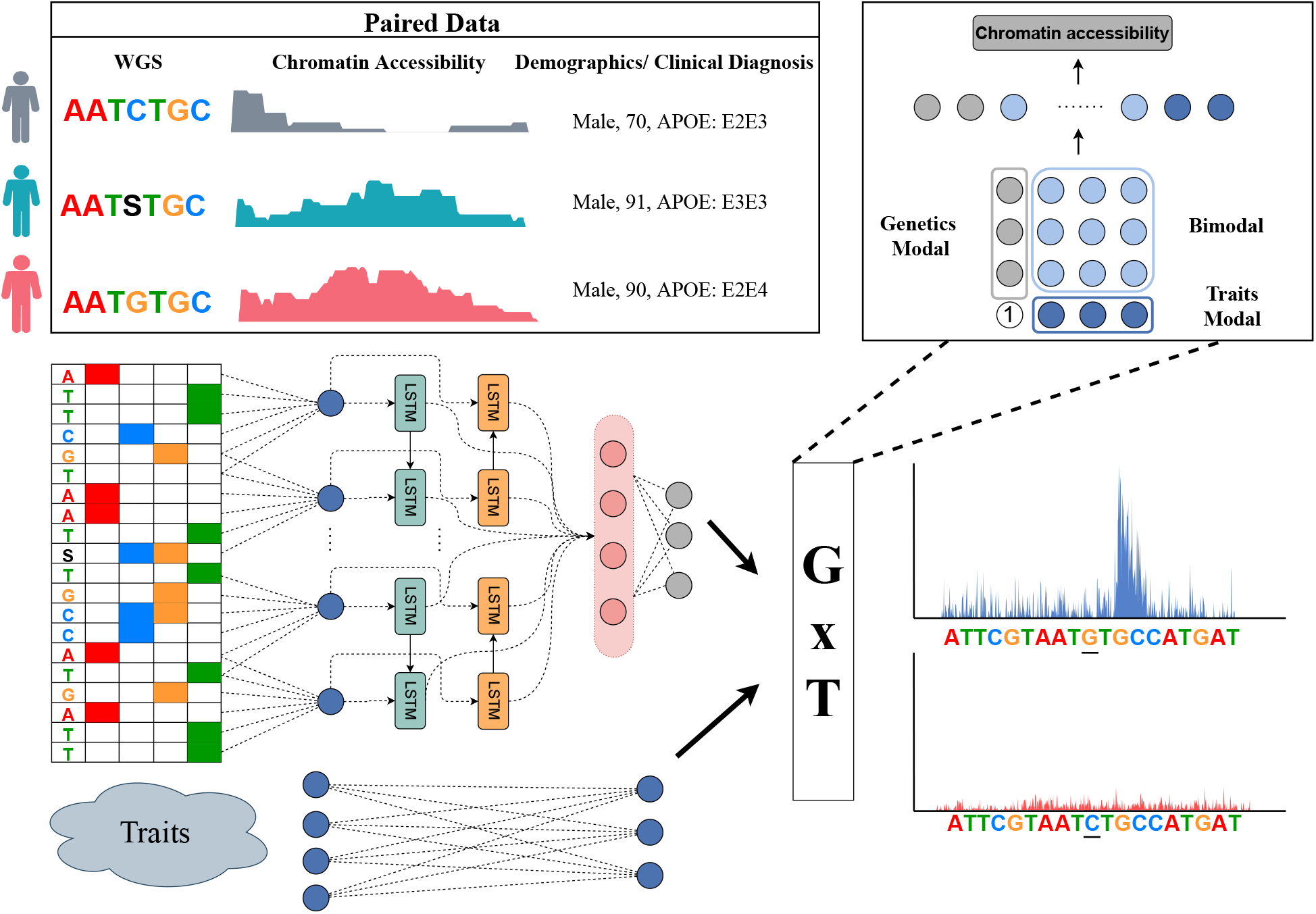
Flowchart of DeepPerVar. DeepPerVar is essentially a multi-modal deep convolutional neural network, which consists of two modalities for handling personal sequence and personal traits as input respectively. DeepPerVar is trained using paired genotype data and epigenetic data in a population study.

**Figure 3:**
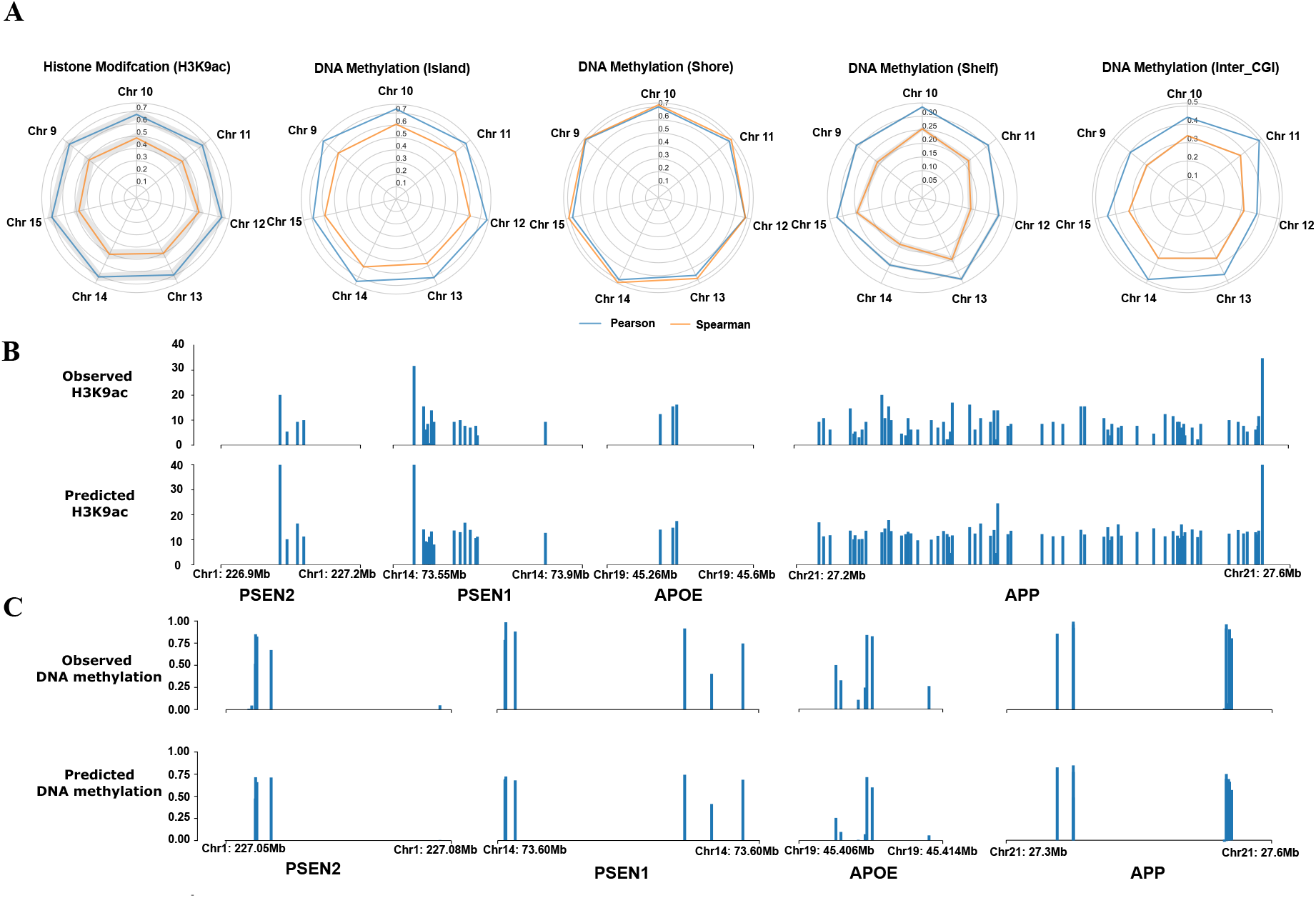
(A) In the radar plot, Pearson correlation (*R*) and Spearman’s rank correlation coefficient (*ρ*) between predicted and observed H3K9ac signals, and methylation ratios in four different DNA methylation regions (island, shelf, shore and open sea) on chromosome 9 to 15 in the testing set. The mean of *R* is represented by the solid blue line, the mean of *ρ* is represented by the solid yellow line, and the standard deviation is represented by the shadow. (B) Visualization of the observed and predicted H3K9ac signals in the neighborhoods of four well-known AD causal genes (PSEN1, PSEN1, APOE, APP). (C) Visualization of the observed and predicted DNA methylation ratio in the neighborhoods of four well-known AD causal genes (PSEN1, PSEN1, APOE, APP)

Since DeepPerVar is trained using genotype and epigenetic data derived from prefrontal cortex of AD patients, we are interested in evaluating whether DeepPerVar can accurately predict epigenetic signals in the key functional genomic sites associated with AD on an individual level. Without loss of generality, we collect one individual (projid=20261037) profiled with H3K9ac and one individual (projid=10415168) profiled with DNA methylation data from ROSMAP. We focus on three causal genes associated with early on-set AD, which include PSEN1 (chr14:73603143-73690399), PSEN2 (chr1:227058273-227083804) and APP (chr21:27252861-27543138) as well as one causal gene APOE (chr19:45409039-45412650) associated with late on-set AD. We plot the predicted normalized reads counts of H3K9ac (Figure 3B) and DNA methylation ratios (Figure 3C) in the neighborhoods of the four genes. We find that DeepPerVar can accurately predict the epigenetic signals in the neighborhoods of the four genes, which indicate that DeepPerVar can potentially be a useful tool to predict epigenetic signals on an individual level, using genotype data only.

### 3.4 DeepPerVar learns known canonical motifs and motifs associated with Alzheimer’s disease

Though deep convolutional neural networks outperform simple linear model in the prediction task by capturing nonlinear dependency among sequence features, they are not readily interpretable as the linear counterparts. However, a common practice to interpret deep convolutional neural network for analyzing biological sequence is to exam the convolutional filters [31]. Specifically, the convolutional filters of the first convolutional layer are usually deemed as the “motif detector”. Each convolutional filter is a 4 × *m* matrix, where *m* is the length of the filter and 4 is the number of nucleotides “A”, “C”, “G” and “T”. Benefiting from the one-hot encoding, the shape of the convolutional filters can mimic the position weight matrix (PWM), which is a widely used representation of motifs in biological sequences. When the convolutional filters scans the DNA sequence in the model training, their weights are updates and learn the pattern of motifs. Therefore, by minimizing the loss function, deep convolutional neural networks can improve the prediction performance for qualitative/quantitative epigenetic signals and capture protein binding information in the epigenetic regions by learning the convolutional filters simultaneously. It should be noted that no prior biological information is used, and all the information can be obtained from the end-to-end training process.

Since H3K9ac play a key role in gene regulation by being an essential part of the active promoter state, we expect that DeepPerVar can learn known canonical motifs as well as detect motifs reported to be associated with AD. By comparing convolutional filters of the first convolutional layer to known human motifs in CIS-BP (See Method section), we find that 455 out of the 512 filters of DeepPerVar-H3K9ac (88.7%) and 372 out of the 512 filters of DeepPerVar-methy (72.7%) are aligned to canonical motifs given a q-value threshold of 0.1. As an example, we plot the sequence logo of five canonical motifs collected from CIS-BP and most matched convolutional filters of DeepPerVar, which include ZNF597 and ZNF749 (zinc finger protein binding motifs), High Mobility Group 20B (HMG20B), Lysine Demethylase 2B (KDM2B) and Transcription Factor 7 (TCF7) (Figure 4A). We find DeepPerVar can accurately discover the patterns of these canonical motifs. In addition, it is reported that several motifs are key functional promoter elements for AD causal genes. For example, part of APP promoter activity is affected by the binding of CCCTC-binding factor (CTCF), stimulating protein 1 (SP1) and upstream stimulatory factor (USF); altering the core sequence region within a binding motif in ETS promoter will results in a drastic decrease in PSEN1 promoter activity; and the proximal C box binds SP1 in APOE promoter and is required for maximum transcriptional activity of APOE [32]. Accordingly, DeepPerVar can identify these AD-associated motif patterns by the convolutional filters (Figure 4B).

**Figure 4:**
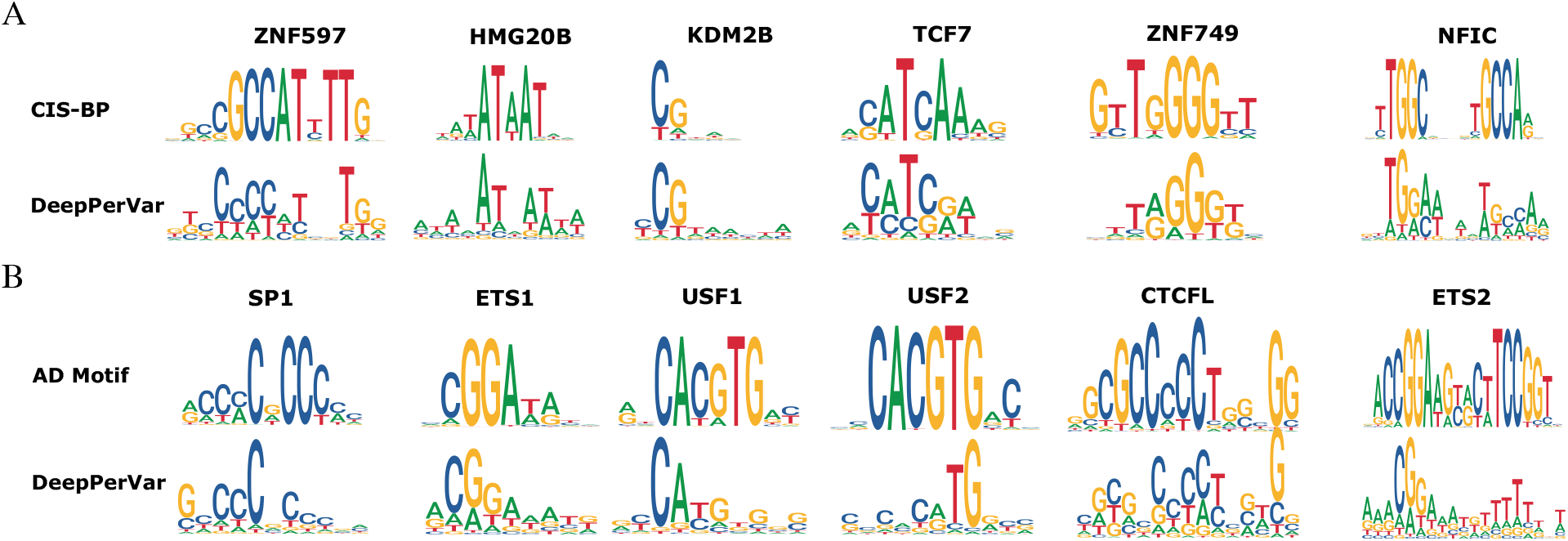
DeepPerVar recovers known canonical motifs and motifs associated with Alzheimer’s disease. (A) Examples of detected discovered motifs with a high similarity to canonical motifs collected from CIS-BP. (B) Examples of discovered motifs with a high similarity to motifs are key functional promoter elements for AD causal genes.

### 3.5 Predicting quantitative epigenetic signals in Roadmap Epigenomics

In the previous section, we have adopted a “cross-sample cross-chromosome” strategy to evaluate the performance of DeepPerVar in predicting H3K9ac signals and DNA methylation ratio in ROSMAP cohort respectively. Here, we will extend the evaluation of DeepPerVar by using a large-scale independent epigenetic datasets in Roadmap Epigenomics Project [8], which has profiled histone modification and DNA methylation across multiple reference tissues and cell types in human. Though DeepPerVar-H3K9ac and DeepPerVar-methy are trained using epigenetic data from prefrontal cortex in brain tissue, we will evaluate whether DeepPerVar has a generalized prediction ability for quantitative epigenetic signals in different tissues and cell lines, which are not included in the model training.

We adopt the same data processing pipeline to obtain the normalized read counts in ChIP-seq peaks as the quantitative signals of 7 core histone modifications, which include H3K4me1, H3K4me3, H3K9ac, H3K27ac, H3K36me3, H3K9me3 and H3K27me3 across 83 cell lines. H3K4me1, H3K4me3, H3K9ac, H3K27ac, H3K36me3 are active/open chromatin marks. Specifically, H3K4me1 and H3K27ac is associated with the high activation of gene transcription and therefore deemed as an active enhancer mark. Similarly, H3K4me3, H3K9ac are defined as active promoter marks and H3K36me3 is active gene body mark. H3K9me3 and H3K27me3 are heterochromatin marks, which are associated with transcriptional repression [33, 34], and therefore are deemed as repressive marks.

We apply DeepPerVar-H3K9ac to predict quantitative signals of 7 histone marks across 83 cell lines/tissues in 18 tissue classes. Pearson correlation *R* between predicted and observed signals is used to evaluate the prediction performance. We visualize *R* for each cell line/tissue and each histone mark in a heatmap and plot the distribution of *R* across all cell lines/tissues for each histone mark (Figure 5A). As a result, H3K4me3 and H3K9ac have best prediction performance (median *R*=0.526 for H3K4me3; median *R*=0.493 for H3K9ac) followed by H3K27ac (median *R*=0.289). It is expected to observe the superior performance of H3K9ac to other histone marks since the training and testing set are from the histone mark though not from the same cell line/tissue. As expected, there is a decline of the prediction performance compared to the “cross-sample cross-chromosome” strategy as the model is evaluated on different cell lines/tissues. Since H3K4me3 and H3K9ac are both active chromatin markers and occur at the promoter of active genes, the prediction performance of H3K4me3 is similar to H3K9ac. In contrast, the prediction performance of heterochromatin marks such as H3K27me3 and H3K9me3 is poor (median *R*=0.218 for H3K27me3; median *R*= 0.018 for H3K9me3), which indicate that active chromatin markers share different regulatory grammar as heterochromatin markers. Interestingly, the performance of active chromatin mark H3K36me3 is also poor mainly because it occurs in body of active genes, which has a different sequence grammar as the promoter. Moreover, H3K4me1, which is an active enhancer mark, also has undesirable prediction performance. This observation indicates that enhancer and promoter may have different regulatory grammar.

**Figure 5:**
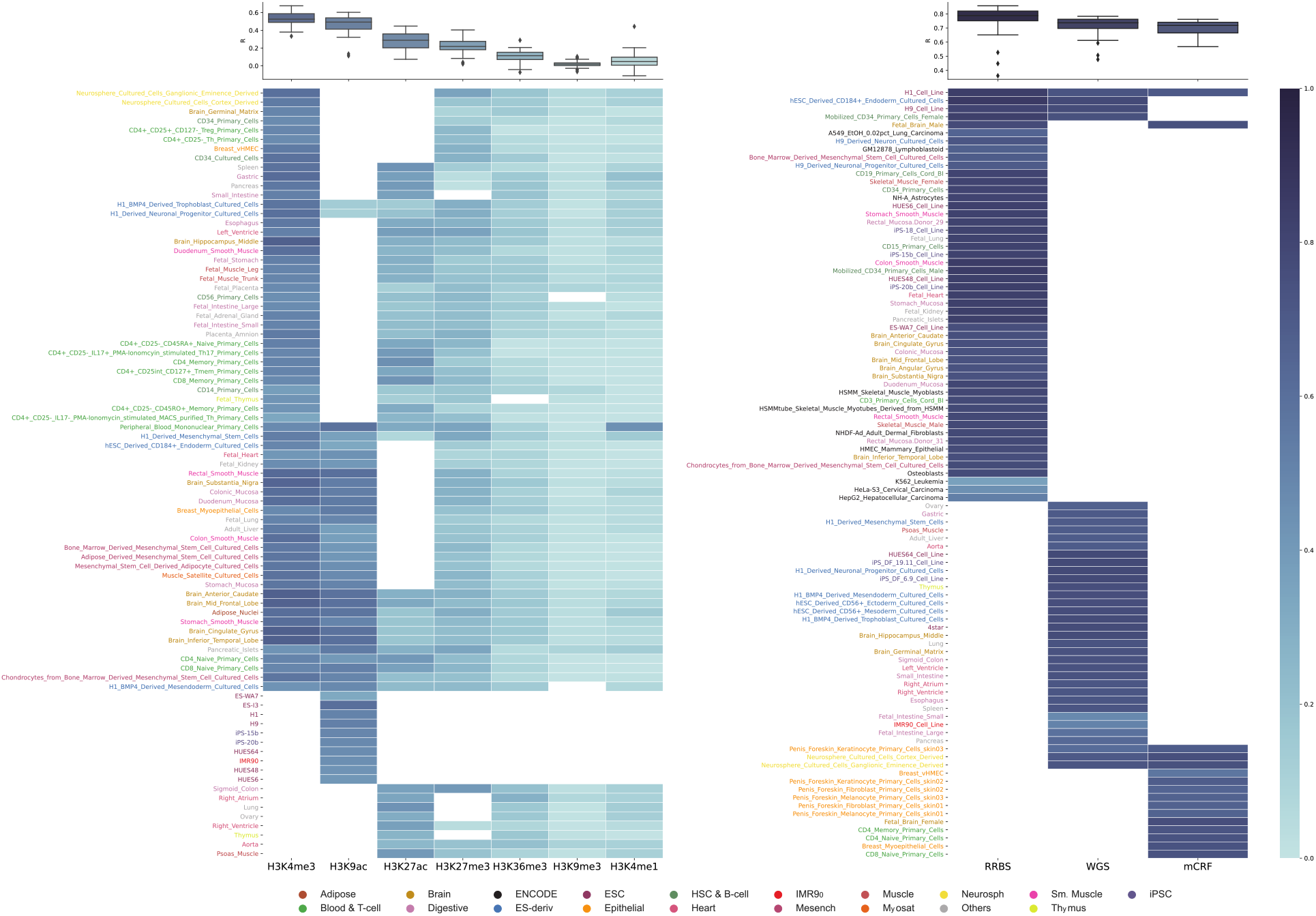
(A) DeepPerVar-H3K9ac to predict quantitative signals of 7 histone marks across 83 cell lines/tissues in 18 tissue classes. (B) DeepPerVar-methy to predict the methylation ratio for datasets generated by three different sequencing techniques (RRBS, WGBS, and mCRF) across 95 cell lines/tissues in 17 tissue classes.

For H3K9ac, we exam *R* is five brain tissues, which include Middle Frontal Lobe (*R*=0.591), Cingulate Gyrus (*R*=0.589), Inferior Temporal Lobe (*R*=0.600), Substantia Nigra (*R*=0.535) and Anterior Caudate (*R*=0.590). Overall, *R* in brain tissues ranges from 0.535 to 0.600 with a median 0.590 compared to *R* ranging from 0.113 to 0.603 with a median 0.477 in other tissues. We further perform a t-test between *R* in brain tissues and *R* in other tissues and find *R* in brain tissues are significantly higher (pvalue=1.829×10^−06^). This observation can be explained by both the training set and testing set are from brain tissues even if not exactly the same as prefrontal cortex, where the training set of DeepPerVar-H3K9ac comes from. Similarly, for H3K4me3, we investigate *R* in seven brain tissues, which include Germinal Matrix (*R*=0.498), Cingulate Gyrus (*R*=0.677), Substantia Nigra (*R*=0.645), Inferior Temporal Lobe (*R*=0.676), Middle Hippocampus (*R*=0.676), Anterior Caudate (*R*=0.671) and Middle Frontal Lobe (*R*=0.667). *R* in brain tissues ranges from 0.498 to 0.677 with a median 0.671 compared to *R* ranging from 0.334 to 0.625 with a median 0.517 in other tissues. A t-test between *R* in brain tissues and *R* in other tissues is performed and *R* in brain tissues are significantly higher than non-brain tissues (pvalue=0.002). However, the difference of *R* between brain and non-brain tissues is not significant for other histone marks (Figure S2).

Moreover, we calculate the DNA methylation ratio for datasets generated by three different sequencing techniques (RRBS, WGBS, and mCRF) across 95 cell lines/tissues in 17 tissue classes. Each sequencing technique targets a subset of cell lines/tissues (Figure 5B). Then, we apply DeepPerVar-methy to predict the methylation ratio and use *R* as the evaluation metric. For RRBS data, *R* ranges from 0.362 to 0.857 (sd=0.095) with a median 0.788 for 51 cell lines/tissues in 13 tissue classes. For WGBS data, *R* ranges from 0.477 to 0.783 (sd=0.071) with a median 0.737 for 37 cell lines/tissues in 13 tissue classes. For mCRF data, *R* ranges from 0.568 to 0.762 (sd=0.051) with a median 0.718 for 16 cell lines/tissues in 5 tissue classes.

Similarly, we plot the heatmap on the level of tissue class for both histone marks and DNA methylation (Figure S1) and find the overall trend is similar to the level of cell lines/tissues. To sum up, we find that DeepPerVar-H3K9ac can accurately predict the epigenetic signals for active chromatin marks associated with the promoter even if the cell lines/tissues of the training set and testing set are different. However, the prediction performance of active chromatin marks associated with the enhancer and gene body, as well as repressive chromatin marks, are undesirable, which indicate these marks share different regulatory grammars as the active promoter marks. Moreover, the prediction performance in brain tissues is overall better than other tissues by benefiting from the both training set and testing set are from the brain region. DeepPerVar-methy achieves an overall stable prediction for methylation ratios across cell lines/tissue and sequencing platforms. Altogether, both evaluations of prediction performance for core histone modifications and DNA methylation in multiple cell lines and tissues indicate that DeepPerVar holds a generalized power to predict epigenetic signals in a broad context.

### 3.6 Comparison between DeepPerVar and state-of-the-art deep learning models in partitioning heritability

DeepSEA and its extension ExPecto, and DeepFIGV are two state-of-the-art deep learning models to evaluate the functional consequence of genetic variants by calculating the allelic difference of epigenetic and transcriptomic signals. Specifically, DeepSEA/ExPecto is a multi-task deep convolutional neural network provide a binary prediction for TF binding event or chromatin accessibility using the universal reference genome only. DeepPerVar is similar to DeepFIGV on modeling paired WGS and epigenetic assays in a population study. However, we want to point out that DeepPerVar differs DeepFIGV in several important aspects. First, the network structure is different. Besides standard convolutional layers to learn local sequence correlation, DeepPerVar adopts Bidirectional LSTM layer to capture long distance dependency for DNA sequence. In addition, DeepPerVar is essentially a multi-modal model, which includes one subnetwork to learn sequence representation and another subnetwork to learn trait representation. Importantly, DeepPerVar develops a novel fusion module layer, which can jointly learn the unimodal representation and bimodal interactions from both sequence and trait representations. Second, DeepFIGV focus on modeling lymphoblastoid cell lines in 75 normal individuals from 1000 Genome Project. However, DeepPerVar focus on a much larger Alzheimer’s disease cohort (∼ 200 individuals) and therefore can unveil potentially genetic mechanism of disease.

All the three deep learning methods can provide genome-wide functional scores to measure the confidence of functional variants. Without loss of generality, we use DeepPerVar-H3K9ac for the partitioning heritability analysis. To compare DeepPerVar and its competitors, we first rank all variants in genome-wide association summary statistics using functional scores from different deep learning methods, and then perform partitioning heritability analysis for top-ranked variants at different top percentages. Since functional categories of the genome contribute disproportionately to the heritability of complex diseases, we expect the top-ranked variants will explain more heritability in functional annotations that are derived from tissues and cell types, which are relevant to the studied complex disease. Accordingly, a superior deep learning method will not only show enrichment for cell types and tissues that are associated with the complex disease but also heritability enrichment of top-ranked variants in key functional elements.

To do this, we collect summary statistics for Alzheimer’s dementia from Iris Jansen et al. [35]. Then, we obtain the functional scores of three deep learning methods for all variants and rank all variants in a decreasing order. For variants at different top percentages (1%, 3%, and 10%) in the rank list from each deep learning method, we adopt stratified LD score regression [36] to estimate polygenic contributions to heritability. Each functional annotation is deemed as a combination between cell type/tissue and regulatory element. The key regulatory element under the consideration include active enhancer marks (e.g., H3K27ac, H3K4me1), active promoter marks (e.g., H3K9ac, H3K4me3), active gene body mark (e.g, H3K36me3) as well as DNase I hypersensitive sites. There are 220 cell type-specific functional annotations mostly derived from Roadmap Epigenomics and ENCODE. We further categorize the 220 cell type-specific annotations into 10 cell type groups in the same way as Finucane et al. [36], which include adrenal or pancreas, CNS, cardiovascular, connective and bone, digestive, immune or blood, kidney, liver, skeletal muscle and other.

For the cell-type-specific analysis (Figure 6A), we find DeepPerVar obtains the highest enrichment in CNS for top 1% SNPs and the enrichment declines when the top percentage increases, which indicates that DeepPerVar can potentially prioritize putative causal variants enriched in cell types relevant to AD. Since CNS is well-known to be associated with AD disease etiology, this observation unveils the underpinning biology. However, compared to DeepPerVar, the enrichment of CNS dramatically declined in DeepFIGV and ExPecto, which indicates that DeepPerVar can better identify cell types relevant to AD by leveraging the paired genotype and epigenetic data in ROSMAP AD study.

**Figure 6:**
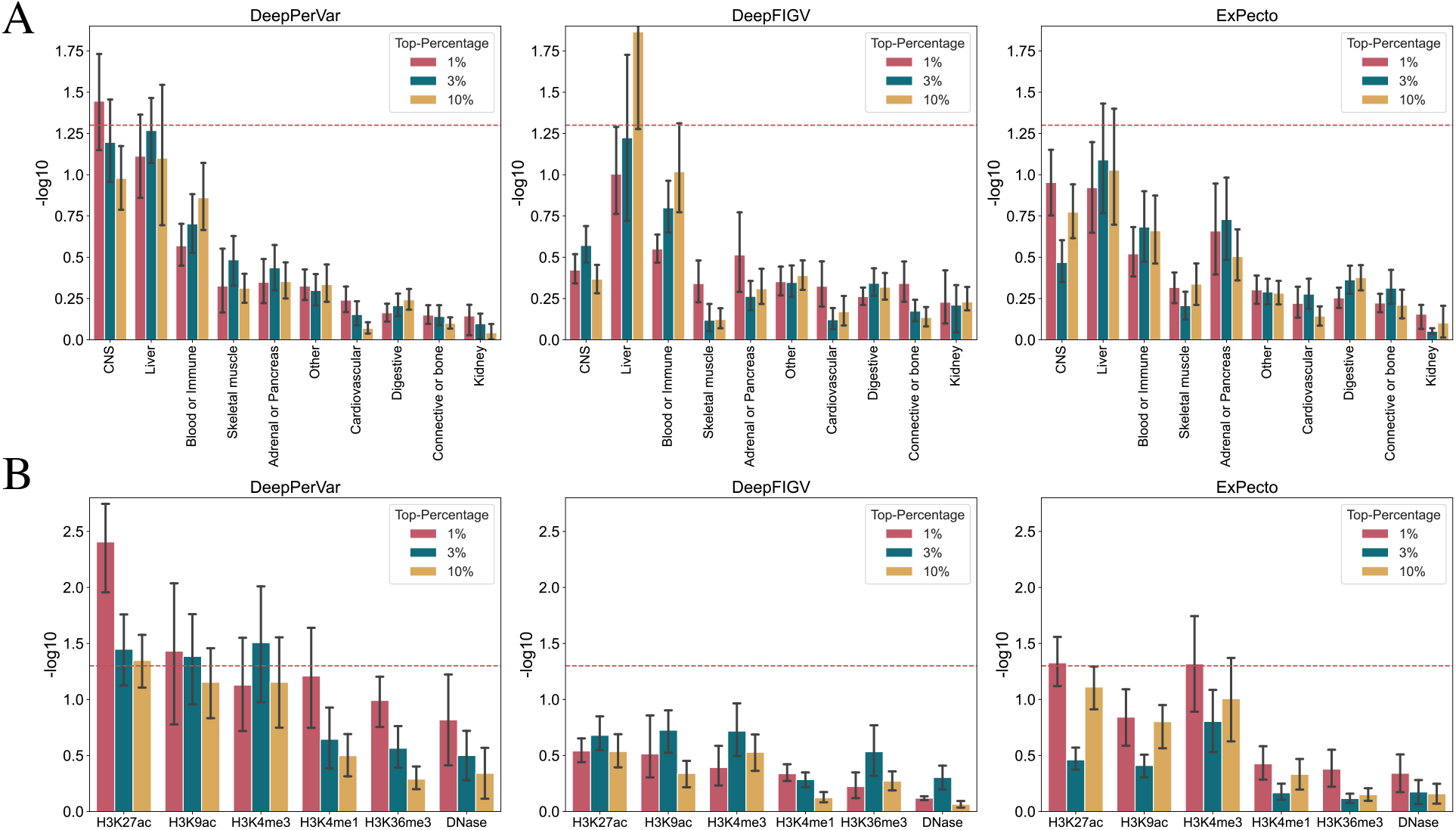
Comparing DeepPerVar to DeepFIGV and ExPecto in partitioning heritability using LD score regression. (A) Cell type-specific analysis for 10 cell groups for top 1%, 3% and 10% variants in one AD summary statistics. (B) Regulatory element-specific analysis for core histone marks and open chromatin for top 1%, 3% and 10% variants in CNS. Error bars indicate 2 standard deviations. Red dash line is −*log*_10_(0.05).

Then, we perform the regulatory element-specific analysis within CNS (Figure 6B). DeepPerVar obtains a significant enrichment for H3K27ac, which indicates H3K27ac contributes significantly to SNP heritability for top 1% SNPs. And the enrichment declines as the top percentage increases. Following H3K27ac, H3K9ac achieves a second highest enrichment. The high enrichment of H3K27ac and H3K9ac is further supported by the finding that enrichment in H3K27ac and H3K9ac are linked to transcription, chromatin and disease pathways in AD [37]. The declined enrichment for H3K27ac and H3K9ac with the increase of top percentage indicates that DeepPerVar can prioritize putative causal variants associated with regulatory elements, which play an important functional role to AD. Moreover, as DeepPerVar is trained using H3K9ac, H3K9ac is expected to achieve a higher enrichment than other regulatory elements such as H3K4me1, H3K36me3 and DNase. It is also interesting to observe a similar performance between H3K4me3 and H3K9ac as both marks are associated with active promoters. Moreover, except H3K4me3, the enrichment of other histone marks declines with the increase of top percentage. Compared to DeepPerVar, the enrichment of all regulatory elements in DeepFIGV and ExPecto declines significantly across all levels of top-ranked SNPs. However, the trend of ExPecto is similar to DeepPerVar, where the enrichment of H3K27ac, H3K9ac and H3K4me3 is significantly higher than other regulatory elements.

### 3.7 Prioritization of putative causal variants associated with Alzheimer’s disease in a risk locus

Typically, GWAS can identify common variants which tag a region of linkage disequilibrium (LD) containing causal variants. However, it is challenging to determine the underlying causal variants by using genotype data only due to the complex patterns of linkage disequilibrium among SNPs. Therefore, the tag SNPs are not necessarily to be causal but the variants in LD can be the causal ones. Here, we adopt a similar statistical measure [38], which is based on the functional scores of DeepPerVar-H3K9ac, to detect noncoding variants that are most likely to be causal to AD.

Moreover, Psychosis (delusions or hallucinations) in Alzheimer’s disease (AD+P) occurs in up to 50% of AD patients, and is associated with significantly worse clinical outcomes. Drug such as Atypical antipsychotics, which is originally developed for Schizophrenia, are also commonly used for the AD+P treatment [39]. In addition, one recent study unveils that there are shared genetic underpinning between AD+P and Schizophrenia by finding Schizophrenia polygenic risk score (PRS) is associated with AD+P [40]. Therefore, prioritization of putative causal variants associated with AD in a GWAS risk locus of Schizophrenia will give additional evidence of the shared genetic liability between AD and Schizophrenia.

To create a risk locus, we first choose a GWAS SNP rs1658810 (pvalue=2×10^−13^), which is a non-coding variant positively associated with Schizophrenia and located in the intron of C2orf69. C2orf69 is a protein-coding gene, which plays an active role in the development/homeostasis of the immune and central nervous systems [41]. Moreover, in the eQTL analysis, rs1658810 is also found to be significantly associated with C2orf47 (pvalue=1.30×10^−5^) and LINC01792 (pvalue=8.29×10^−4^) in CommonMind [42]; associated with TYW5 in both GTEx [43] (pvalue=1×10^−6^ for dorsolateral prefrontal cortex) and Common-Mind (pvalue=2.25×10^−9^); and associated with FTCDNL1 in GTEx (pvalue=8.30×10^−6^ for dorsolateral prefrontal cortex) and CommonMind (pvalue=2.97×10^−13^). Besides the evidence from both GWAS and QTL analysis, a recent functional study using massively parallel reporter assay (MPRA) further validates that rs1658810 shows allelic differences in driving reporter gene expression into both K562 chronic myelogenous leukemia lymphoblasts and SK-SY5Y human neuroblastoma cells [44]. All these experimental and computational evidences indicate that neighborhood region of rs1658810 is a highly confident risk locus.

Specifically, we extend 100kb upstream and downstream of rs1658810 as the GWAS risk locus and curate all noncoding variants from 202 AD individuals in ROSMAP in the risk locus, which results a total of 2159 variants. To demonstrate the prioritized variants are more likely to be AD causal, we assess the relationship between prioritized variants and Braak staging, which is a quantitative trait to classify the degree of pathology in AD. First, we filter out the variants with minor allele frequency less than 10 to ensure the number of individuals is enough for evaluation, yielding 511 variants. Second, for each variant, we calculate the allelic difference of Braak staging by computing the absolute value of the change between the average Braak staging of individuals with the minor alleles and those with the major alleles. Last, we plot the allelic difference of average Braak staging for K top-ranked variants versus K bottom-ranked variants, ordered by functional score of DeepPerVar-H3K9ac (Figure 7). As a result, we find that top-ranked putative causal variants are more highly correlated with change of Braak staging than bottom-ranked ones. When K equals 20 and 40, the difference between change of Braak staging between top and bottom variants are statistically significant (pvalue*<*0.05). However, the significance diminishes when K increases. Therefore, by checking the correlation between the genotypes of variants and Braak staging, we demonstrate that DeepPerVar-H3K9ac is effective in the prioritization of putative causal variants in a GWAS risk loci of Schizophrenia. In addition, this finding strengthens the evidence of shared genetic liability between AD and Schizophrenia.

**Figure 7:**
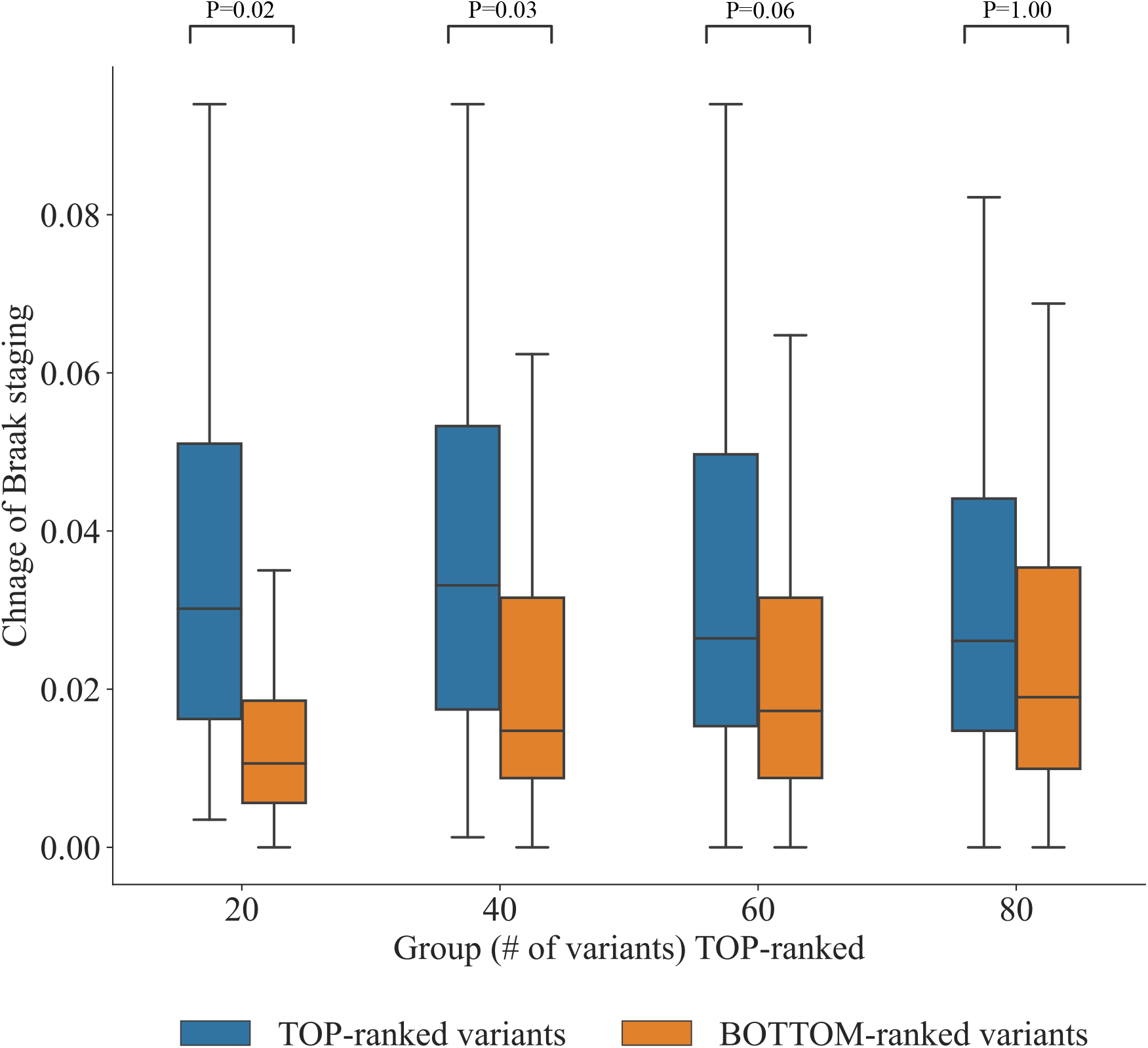
Functional scores of DeepPerVar-H3K9ac can help prioritize putative causal variants in a GWAS risk locus associated with Schizophrenia. The risk locus is defined as 100kb upstream and downstream of GWAS SNP rs1658810 associated with Schizophrenia. All variants in the risk locus are ranked by the functional scores in a decreasing order. For each variant, we calculate the allelic difference of Braak staging, which is absolute value of the change between the average Braak staging of individuals with the minor alleles and those with the major alleles from AD individuals in ROSMAP. The allelic difference of Braak staging of K top-ranked putative causal variants and K bottom-ranked variants (K = 20, 40, 60, and 80) are plotted.

## 4 Discussion

To identify and interpret the functional consequence of genetic variants, especially the noncoding ones, is a challenge in genetics research. GWAS can identify common variants associated with complex diseases. However, these GWAS SNPs may not be the causal due to linkage disequilibrium and limited statistical power. Moreover, WGS analysis can identify additional rare variants. Nevertheless, there is a lack of fine resolution for state-of-the-art statistical approaches to pinpoint the casual variants in the association region. In addition, QTL analysis can identify variants associated with molecular traits such as gene expression or epigenetic signals. Yet, these molecular trait QTL analysis is subject to the same challenges of linkage disequilibrium and limited statistical power as GWAS/WGS so they cannot pinpoint causal variants either. Moreover, all these association analysis cannot predict the functional consequence of a variant not observed in the analyzed genotype data.

To provide tissue/cell type-specific interpretation for evaluating how noncoding variants affect molecular mechanisms to have a functional impact on the disease, enrichment analysis becomes a popular way to annotate variants with GWAS risk variants by positional overlap with tissue/cell type-specific functional annotations. In addition, using labelled causal variants (e.g, HGMD and ClinVar) with functional annotations, multiple supervised machine learning models have been developed to achieve a genome-wide prediction by assigning a functional score to each variant. However, these annotation-based approaches do offer an option for evaluating allelic effect of a variant, which is essential to decipher the functional impact of a variant.

Most recently, deep convolutional neural networks have been developed to predict genome-wide transcription factor binding, histone modification and chromatin accessibility across thousands of tissues and cell types in a multi-task learning framework using DNA sequence. These approaches are flexible to *in silico* evaluate allelic effect of a variant by comparing the difference of predicted epigenetic signals between reference allele and alternative allele in the reference sequence. Since these methods are developed on sequence context with base-level resolution, they are less susceptible to linkage disequilibrium and can predict the functional consequence of *de novo* variants, which may not exist in the observed genotype data. Since these approaches use reference genome and epigenetic landscapes from a diverse of tissue and cell types, they may be suboptimal in a population study, where heterogeneity of genetic variants and epigenetic landscape on an individual level should be considered. A single-task deep convolutional neural network, named DeepFIGV, has been recently developed to perform quantitative epigenetic variation in chromatin accessibility and histone modifications in a population study of paired WGS and ChIP-seq data for lymphoblastoid cell lines. However, DeepFIGV lacks the disease focus and the consideration of personal traits which are important factors affecting epigenetic signals in a population study.

In this work, we introduce a multi-modal deep convolutional neural network, named DeepPerVar, to characterize the functional effect of noncoding variants in a personal genome in a study population with the disease focus. Since DeepPerVar is single-task model, which targets one specific disease, it might be more optimal in deciphering functional consequence of genetic variants in disease etiology. Different from reference genome-based approaches such as DeepSEA/ExPecto, DeepPerVar models the the contribution of personal genetic variations on epigenetic signals by leveraging paired WGS data and epigenetic data in a population study. Compared to DeepFIGV, DeepPerVar, includes personal traits such as age and gender, which are important factors associated with epigenetic change, and adopt a novel fusion layer to integrate feature representations from both personal genome and personal traits in the model training. To validate this hypothesis, we first demonstrate that both genetic variations and traits affect epigenetic signals on an individual level by real data exploration.

To demonstrate the usefulness of DeepPerVar, we train two versions of DeepPerVar, named DeepPerVar-H3K9ac and DeepPerVar-methy using paired WGS and H3K9ac ChIP-seq, and WGS and DNA methylation data from an AD cohort-ROSMAP, respectively. To evaluate the prediction performance for epigenetic signals, we adopt a conservative “cross-sample cross-chromosome” strategy to create the training, validation, and testing sets from different chromosomes in different individuals. As a result, DeepPerVar obtains an accurate prediction for both H3K9ac signals and DNA methylation ratios. Moreover, DeepPerVar accurately predict epigenetic signals in the key functional genomic sites associated with AD such as early on-set AD casual genes (i.e., PSEN1, PSEN1 and APP) and late on-set AD (i.e., APOE) on an individual level. In addition, DeepPerVar can learn canonical motifs and motifs reported to regulate expression of AD causal genes. Furthermore, we apply DeepPerVar to predict quantitative epigenetic signals in multiple cell lines and tissues in Roadmap Epigenomics Project to independently evaluate whether DeepPerVar has a generalized prediction ability for quantitative epigenetic signals in different cell lines and tissues, which are not included in the model training. We find that DeepPerVar-H3K9ac achieves accurate prediction of H3K4me3 and H3K9ac signals, especially in brain tissues. Interestingly, we DeepPerVar-methy obtains a consistently accurate prediction for DNA methylation ratios even the testing cell lines/tissues are different from the training set-prefrontal cortex.

We perform two experiments to demonstrate that DeepPerVar can interpret and prioritize putative causal variants. First, we rank all variants by DeepPerVar functional scores in one AD GWAS summary statistics and we adopt stratified LD score regression to estimate polygenic contributions to heritability for top-ranked variants with different top percentage. For the cell-type-specific analysis, CNS has the highest enrichment for top 1% SNPs and the enrichment declines when the top percentage increases, which indicates that Deep-PerVar can prioritize putative casual variants enriched in cell types relevant to AD. Moreover, the enrichment of CNS is significantly higher in DeepPerVar than DeepFIGV and ExPecto, which indicates that DeepPerVar can better identify cell types that are mostly relevant to AD. For the regulatory element-specific analysis within CNS, we find the active chromatin marks H3K27ac and H3K9ac are more enriched than other regulatory elements, which indicates their important roles in AD. The enrichment of most regulatory elements also declined with the increase top percentage, which indicates that DeepPerVar can prioritize putative casual variants enriched in regulatory elements that play an important functional role to AD. Moreover, DeepPerVar achieves an overall higher enrichment across all regulatory elements than DeepFIGV and ExPecto. Second, using DeepPerVar functional score, we rank all variants in a highly confident risk locus of GWAS SNP rs1658810 associated with Schizophrenia, whose functional role is further supported by eQTL analysis and MPRA experiments. We find that top-ranked variants are more highly correlated with one quantitative AD trait-Braak staging than bottom-ranked variants. This finding indicates that DeepPerVar is effective in the prioritization of putative causal variants in a GWAS risk locus. Moreover, this finding strengthens the evidence of shared genetic liability between AD and Schizophrenia.

To sum up, by leveraging paired WGS data and epigenetic data in a population cohort, DeepPerVar is powerful to quantitatively predict genome-wide epigenetic signals and evaluate the functional consequence of noncoding variants by quantifying their allelic effect on the prediction on an individual level. In this way, DeepPerVar is capable for the prioritization of putative causal variants in the risk locus and interpretation of noncoding variants in the regulatory process and related to disease etiology. Therefore, DeepPerVar can be a complementary tool to GWAS and QTL analysis, which typically require a large sample size to make sound scientific findings. With the advent and popularity of individual-level multi-omics in a population study such as UK Biobank [54] and NHLBI TOPMed [55], we believe DeepPerVar will be a useful tool in genetics research community.

In this work, we train DeepPerVar using “definite” AD individuals in ROSMAP, so the functional score will measure the functional consequence of a variant on AD disease etiology. We can further extend DeepPerVar to include normal samples as baseline information with an additional personal trait “AD/control” trait subnetwork in the model architecture. Thus, the functional score can be refined by taking the difference between allelic difference of predicted epigenetic signals in AD and control sample. Moreover, we demonstrate the application of DeepPerVar in ROSMAP, which is a cohort profiled with individual-level genotype data and epigenetic data for studying AD. It should be noted that DeepPerVar can be easily adapted to other complex disease, upon availability of paired genotype and epigenetic data, to produce disease-specific functional scores for prioritizing and interpreting putative causal variants.

Notably, DeepPerVar is a “global” model, which utilized all sequence contexts in epigenetic regions across all individuals in the study population. The “global” model, which benefits from training all loci across genome and individuals, is more powerful when the sample size in the study cohort is small. However, when the sample size is large in the study cohort, we can customize DeepPerVar into a region-specific “local” model by using sequence context in the region of interest only. The “local” model will consider the region heterogeneity as well as the population heterogeneity, which can be potentially more powerful in prioritizing and interpreting genetic variants in personal genome. We will focus this direction in the future work.

## Acknowledgement

This work was supported by Indiana University Precision Health Initiative, Showalter Research Trust and National Institute of General Medical Sciences of the National Institutes of Health under Award Number R35GM142701 to LC.

