## Supplemental File for "DeepPerVar: a multimodal deep learning framework for functional interpretation of genetic variants in personal genome"

Ye Wang[1,2], Li Chen[2,3*]

[1] Biotherapeutic and Medicinal Sciences, Biogen, 225 Binney Street, Cambridge, MA, 02142, USA

[2]Department of Biostatistics and Health Data Science, Indiana University School of Medicine, Indianapolis, IN, 46033

[3]Center for Computational Biology and Bioinformatics, Indiana University School of Medicine, Indianapolis, IN, 46033

Table S1: Distribution of phenotypic data for "definite" AD individuals from ROSMAP

| **Variable** | **ROSMAP** |
|---|---|
| Counts (by region) | N=212 |
| Age of dealth (capped at 90) | 87.18 +- 3.75 |
| Sex (% male) | 27.36 |
| Years of education | 16.73 +- 3.47 |
| APOE genotype (% $\epsilon$ 2$\epsilon$ 2/$\epsilon$ 2$\epsilon$ 3/$\epsilon$ 2$\epsilon$ 4/$\epsilon$ 3$\epsilon$ 3/$\epsilon$ 3$\epsilon$ 4/$\epsilon$ 4$\epsilon$ 4) | 0.0/5.6/2.8/51.4/37.2/2.9 |
| CERAD Score | 4.36 +- 0.93 (0.00-3.00) |
| Braak Staging | 3.45 +- 1.22 (0.00-6.00) |
| Clinical cognitive diagnosis(% NCI/MCI/MCI+/AD/AD+/Other) | 13.2/14.1/0.0/63.2/8.0/1.4 |

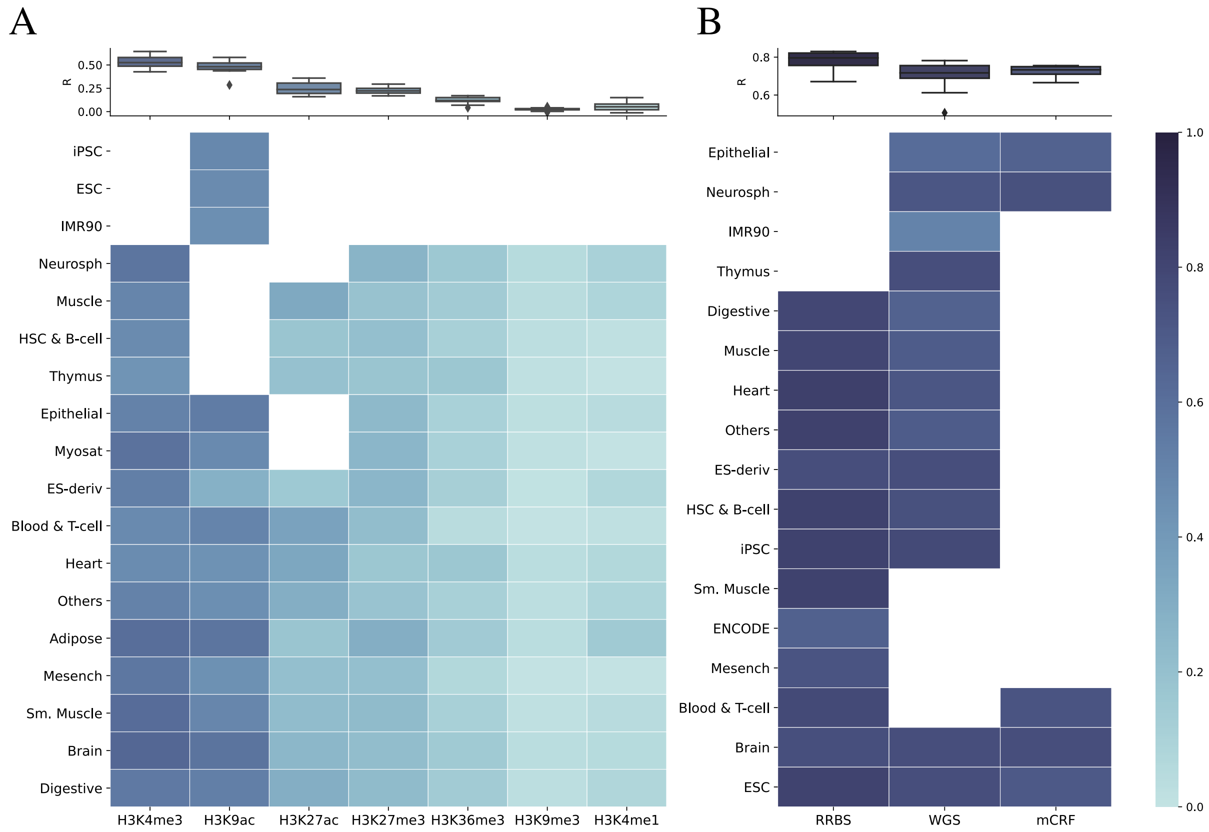


Figure S1: (A) DeepPerVar-H3K9ac to predict quantitative signals of 7 histone marks in 18 tissue classes. (B) DeepPerVar-methy to predict the methylation ratio for datasets generated by three different sequencing techniques (RRBS, WGBS, and mCRF) in 17 tissue classes.

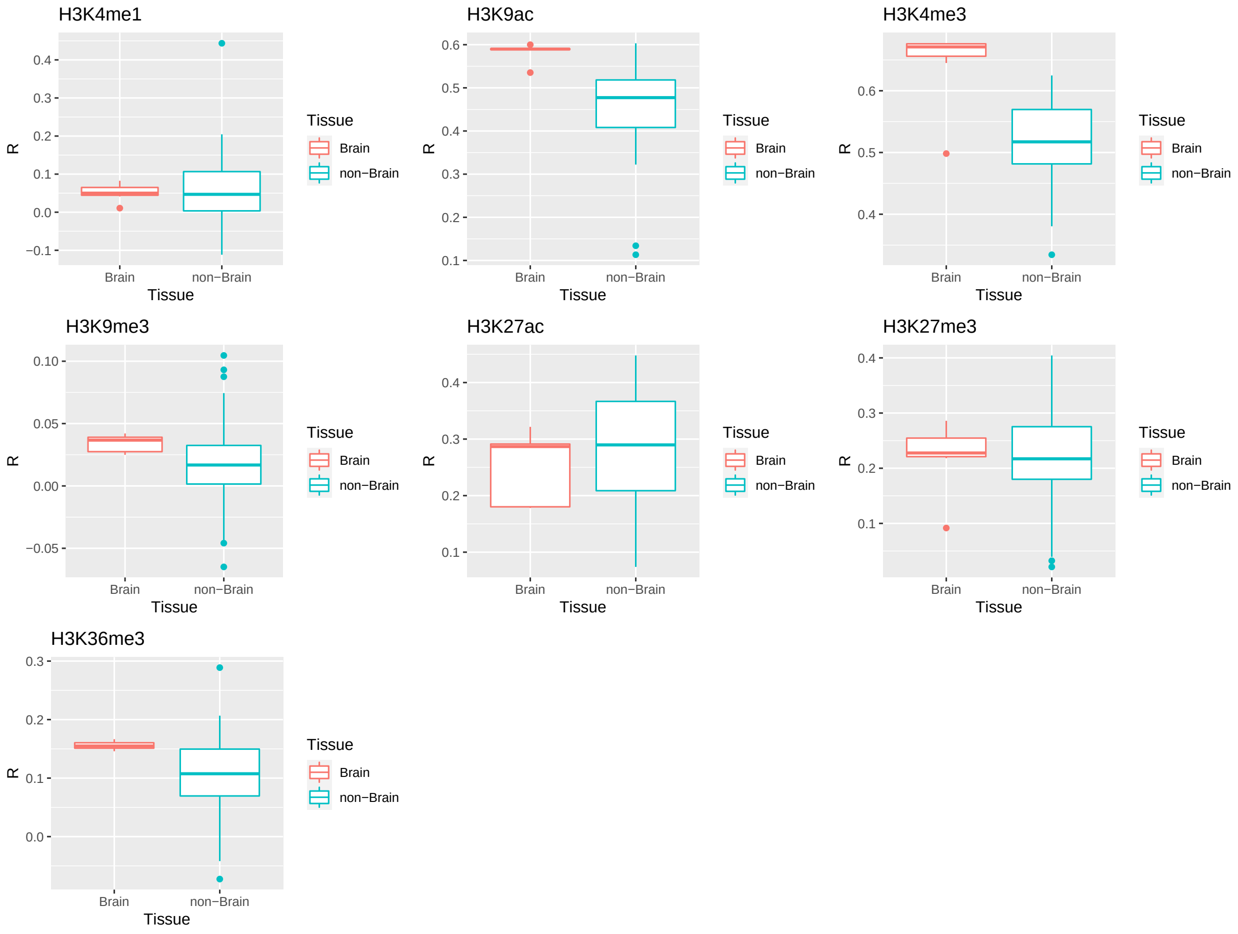


Figure S2: Predicted H3K9ac signals of 7 histone marks between brain and non-brain tissues.
